# Discovering disease genetic variation impacting gene expression in 103 brain tissues with the Brain Ontology Expression (BRONTE) graph neural network model

**DOI:** 10.1101/2025.05.08.652857

**Authors:** Jianfeng Ke, Tingjian Ge, Rachel D. Melamed

## Abstract

Uncovering genes dysregulated in neuropsychiatric diseases promises to put forward therapeutic targets. By linking disease risk variants to tissue-specific gene regulation, the Genotype-Tissue Expression (GTEx) project has enabled discovery of disease genes and the brain tissues where these pathogenic effects occur. But, GTEx requires huge investment in obtaining human brain tissues, limiting the sample size and tissue diversity available. Crucially, this resource does not contain all key tissues where brain diseases are known to originate. Here, we propose a novel graph convolutional model that integrates the GTEx brain expression data with the Allen Brain Atlas, a resource spanning over 100 fine grained brain tissues that are annotated with an ontology of tissue organization. Our model, Brain Ontology Expression (BRONTE), allows completion of the GTEx data for 221 individuals to impute expanded expression in 103 uncollected detailed brain tissues. We show that expression data imputed by BRONTE recovers measured variation in expression across subjects, genes, and brain tissues. Most importantly, our method both recapitulates known relationships between genetic variations and gene expression, and simultaneously discovers both new genes impacted by genetic variants, and the tissues impacted. This allows us to put forward novel mechanisms of effect for genetic variation associated with 30 brain-related traits, supplying therapeutic targets in overlooked tissues of the brain.

## Introduction

Genome-wide association studies (GWAS) of neurological and psychiatric diseases aim to provide insights into mechanisms of disease development and ultimately prioritize therapeutic targets[1], [2]. Therefore, a key goal in analysis of GWAS is to link risk variants to the genes they impact, and the tissues where alterations to those genes drive disease. To enable this linkage, the Genotype Tissue Expression (GTEx) project has comprehensively profiled matched genotype and gene expression across dozens of body tissues from nearly 1000 individuals[3]. The GTEx tissues include around a dozen distinct brain tissues, and these have been used to suggest mechanisms of pathogenicity for variants associated with neurodegenerative[4], [5] and psychiatric[6] disease.

But, the insights enabled by GTEx are limited by the sample size and included tissues of the GTEx data. Large sample size enables discovery of variant effects[3], [7], but most GTEx subjects do not have expression data for all GTEx tissues, and only a few dozen individuals have complete expression data across the GTEx brain tissues. In addition to the number of samples, the diversity of brain samples also limits discovery. The small number of GTEx brain tissues do not represent the exquisite complexity of brain function. For example, initiation of neurodegeneration has been tied to certain cortical tissues not profiled in GTEx[8], [9]. Previous studies have attempted to compensate for this deficit in tissue representation via post-hoc analyses[4], [10]. Instead, we propose that expanding the GTEx data to represent a greater number of individuals for each tissue, and a greater diversity of tissues for each individual, could have great impact for discovering mechanisms of disease and therapeutic targets.

To increase sample size, some previous work has sought to impute missing GTEx tissues[11], [12], [13], but no previous method leverages existing data on brain biology. Here, we create a model incorporating both GTEx and the Allen brain expression atlas, which collected gene expression for over 100 fine-grained brain tissues[14]. The Allen project showed that gene expression follows a tissue-specific expression pattern that is stable across individuals, and that expression is related to an ontology describing the hierarchical relationship of brain tissues. Our model, Brain Ontology Expression (BRONTE), learns from these gene and tissue patterns to predict gene expression in GTEx subjects across 103 Allen brain tissues, including all GTEx tissues. We demonstrate that BRONTE is able to predict variation in gene expression across held out GTEx individuals, and we show that the imputed GTEx expression profiles resemble data from the Allen brain project. Most importantly, the predicted expression data for GTEx subjects enables both recovery of known variant functions, and discovery of new gene dysregulation associated with disease. We expect that this method, and the imputed data set we provide, will expand discovery of genetic variation impacting brain disease.

## Methods

### Data curation

We download transcriptomic data (RNA-seq) of the human brain from the GTEx Portal and the Allen Brain Atlas. The Allen dataset comprises microarray data from 6 subjects, which undergo quantile normalization per gene across all brain tissues and subjects. We then focus on 103 shared tissues within the left hemispheres of these subjects, and average expression values from microarray probes associated with the same gene. For the GTEx dataset, we select 10 out of the 13 available brain tissues, including Amygdala, Anterior cingulate cortex BA24, Caudate basal ganglia, Cerebellar Hemisphere, Frontal Cortex BA9, Hippocampus, Hypothalamus, Nucleus accumbens basal ganglia, Putamen basal ganglia, and Substantia nigra. We exclude three tissues from our analysis: two duplicated tissues, Cerebellum and Cortex, and one tissue absent in the Allen dataset, Spinal cord cervical c-1. Our analysis centers on a common set of 15,044 genes present in both the GTEx and Allen datasets. Mapping between 10 GTEx tissues and 103 Allen tissues and the hierarchical structures of brain tissues are provided in Supplementary Table 1.

**Table 1.** Comparison between BRONTE discovered genes and Metabrain study discovered genes in MR+coloc.

| Trait | BRONTE discovered genes | Metabrain discovered genes | Overlapped genes | P-value |
| --- | --- | --- | --- | --- |
| Cognitive function | 39 | 28 | 11 | 6.84e-19 |
| Years of schooling | 57 | 55 | 10 | 9.13e-14 |
| Schizophrenia | 26 | 27 | 6 | 1.99e-11 |
| MS | 14 | 22 | 5 | 3.03e-11 |
| ALS | 2 | 3 | 1 | 5.19e-04 |
| PD | 3 | 4 | 1 | 1.04e-03 |
| BD | 6 | 4 | 1 | 2.07e-03 |
| AD | 4 | 8 | 1 | 2.77e-03 |
| Putamen volume | 1 | 0 | 0 | 1.00e+00 |
| ADHD | 2 | 0 | 0 | 1.00e+00 |
| Generalized EPI 2 | 3 | 0 | 0 | 1.00e+00 |
| JME | 0 | 1 | 0 | 1.00e+00 |
| MDD | 5 | 2 | 0 | 1.00e+00 |

### Model design and training

To predict the expression of gene g in Allen tissue t of subject s, BRONTE incorporates three components as inputs: learned tissue embeddings and gene embeddings representing tissue t and gene g (described below), and the gene expression vector of gene g from 10 GTEx tissues for subject s (missing tissues can be imputed), representing subject- and gene-specific variation. These inputs are concatenated and fed into a multi-layer perceptron (MLP) architecture with two sets of hidden layers to generate the prediction. Each hidden layer in the MLP consists of a fully connected linear layer followed by a Rectified Linear Unit (ReLU) activation function. To predict the gene expression of a GTEx tissue for Allen subjects, we generate the prediction by averaging the predicted gene expression of the subordinate Allen tissues corresponding to that GTEx tissue, using the Allen ontology. BRONTE is initially trained on 6 Allen subjects and then fine-tuned on 30 GTEx subjects with all 10 GTEx tissues.

#### Tissue Embedding Training

The tissue embeddings are derived from a Graph Convolutional Neural Network (GCN), trained on a graph where nodes represent tissues and edges the hierarchical relationships between tissues. This tree graph is created from the hierarchical ontology of brain tissues provided by the Allen brain project, with the 103 most fine-grained Allen tissues as the tree’s leaves. Each tissue node in the graph is associated with a feature vector, which is initialized to a one-hot, and trained to learn tissue embeddings through model training.

#### Gene Embedding Pretraining

The gene embeddings are intended to represent variation in gene expression across tissues, as described in the Allen brain atlas project. Therefore, we train gene embeddings on the Allen dataset such that the Pearson correlation between gene embeddings is close to the Pearson correlation of gene expression values across the 103 Allen tissues for any pair of genes, for any individual. Training examples are drawn iteratively across each of the 6 Allen subjects.

### Missing GTEx tissue Imputation with BRONTE

We develop 10 extended BRONTEs, one for each GTEx tissue, to impute missing tissues in GTEx subjects. The Allen brain tissues are more fine-grained than the GTEx tissues, with each GTEx tissue represented by one or more Allen brain tissue. To impute a given GTEx tissue, the extended BRONTE uses gene expression data from the other 9 GTEx tissues as the subject gene expression vector s, to generate model predictions for Allen brain tissues subordinate to the GTEx tissue to be imputed. These subordinate tissue gene expression values are then averaged. For subjects with more than 1 missing tissue, a two-step imputation process is applied. For each missing tissue, we first impute other missing tissues with linear regression models trained on known tissues from 30 GTEx subjects with complete data. Then, the respective extended BRONTE is used to predict the missing tissue.

### Model comparison with 4 other models and their implementation

We compare our model with four alternative approaches on the collected GTEx data.

Linear regression models are trained on 30 GTEx subjects with all 10 GTEx tissues. For each subject, we leave out one known tissue at a time and predict the expression in the omitted tissue using data from the remaining tissues.

MICE (Multivariate Imputation by Chained Equations)[15], an R package for multiple imputations (version 3.16.0), follows the same leave-one-out evaluation strategy as the CP method (below). For each subject, we mask one collected tissue at a time and use MICE to predict the expression in the masked tissue. The model is applied across 10 tissues, all genes, and all GTEx subjects with at least 5 tissues, using default parameters.

The CP method[16], implemented via the CANDECOMP/PARAFAC (CP) algorithm in the tensorBF R package (version 1.0.2), is run separately for each collected tissue for each subject. For each prediction, we apply the CP method across 10 tissues, all genes, and all GTEx subjects with data available for at least 5 tissues, using parameters K = 20, prop = 0.5, and conf = 0.5.

PrediXcan[17], a computational tool for predicting gene expression from genotype data, is also included for comparison. We perform predictions on overlapping genes and subjects for each tissue using genotype data downloaded from the GTEx portal.

### Comparison of imputed expression to signal in Allen brain

Differential stability (DS score) was developed in the Allen brain atlas project to measure how consistently a gene is expressed more in one brain tissue than another across all sampled brains, providing a way to quantify the reproducibility of tissue expression patterns[18]. The Allen brain atlas defined DS score as the average Pearson correlation coefficient calculated across all pairs of individuals, where each coefficient represents the correlation of the gene’s expression levels across 103 Allen tissues for a given pair. We compare the DS score of measured expression data against that in our imputed data.

WGCNA is a method used to identify clusters of genes with similar expression patterns[19]. In the Allen brain atlas, this method was used to identify modules of expression across tissues; these modules were largely reproducible across individuals. We apply WGCNA to the gene expression data from 103 predicted Allen tissues for each GTEx subject using WGCNA version 1.72.5. The softPower parameter is set to 14, and default settings are used except for the following adjustments: cutHeight = 0.999, deepSplit = 4, pamRespectsDendro = FALSE, and minClusterSize = 30. Then, we compare the modules learned on our imputed data against the original Allen modules, focusing on the overlapping genes between the two datasets, using the Adjusted Rand Index (ARI). To assess statistical significance, the modules from our study are randomly shuffled, and the ARI is recalculated. This randomization process is repeated 1,000 times to generate a distribution of random ARIs. Finally, we compare the ARI from our results to the distribution of shuffled ARIs to create a direct permutation based p-value.

### eQTL, MR, and colocalization analysis

We perform cis-eQTL analysis on GTEx data using the eQTL discovery pipeline for the GTEx Consortium introduced by Broad Institute[20]. We use Mendelian Randomization (MR) analysis with the Wald ratio method to estimate the causal effect of gene expression (exposure) on brain-related traits (outcome). This approach calculates the change in risk for brain-related traits per standard deviation increase in gene expression influenced by the eQTL. MR analysis is conducted on eQTLs identified as significant through FastQTL permutation-based testing across 103 Allen brain tissues and 30 neurological traits. To enable the LD clumping and GWAS statistic querying steps, we first use the R package SNPlocs.Hsapiens.dbSNP144.GRCh38 (version 0.99.20) to identify the rsid mappings for our SNPs. The eQTLs are then LD-clumped using the ld_clump() function from the R package ieugwasr (version 1.0.1)[21] with default parameters (10,000 kb clumping window with *r*^2^ cut-off of 0.001 using the 1000 Genomes EUR reference panel). GWAS statistics for the eQTLs are retrieved from the GWAS outcomes available in the IEU OpenGWAS dataset using the query_gwas() function in the R package gwasvcf (version 0.1.2)[22]. If a SNP is not available in the outcome GWAS, proxy SNPs are identified through a proxy search with a minimum *r*^2^ threshold of 0.8 relative to the eQTL. Next, we use the harmonise_data() function in the R package TwoSampleMR (version 0.6.8)[23] to align the orientation of effect alleles between the eQTL and outcome associations, selecting Action 1, which assumes all alleles are coded on the forward strand and does not attempt to flip alleles. Finally, we perform MR analysis using mr_singlesnp() in TwoSampleMR. We report all MR findings that remain significant after FDR correction across 11,841 SNPs (the union of all the SNPs across all tissues and traits), 103 Allen brain tissues, and 30 brain-related traits.

To evaluate the causal relationship suggested by MR, co-localization analysis is performed on MR results that pass the FDR correction, assessing whether the eQTL and brain-related traits share a common causal signal. LiftOver[24] is employed to convert the hg19 coordinates for all SNPs in the eQTL dataset to ensure consistency across reference genomes. The genomic region for colocalization is defined by including all SNPs within a 100 kb window centered on the primary SNP of interest, based on hg19 positions. Neighboring SNPs with mismatched chromosome annotations between hg19 and hg38 are excluded to maintain alignment in genomic coordinates. Co-localization analysis is performed using coloc.abf() from the R package coloc[25], employing two parameter configurations: the default setting (p1=p2= 10^−4^, p12= 10^−5^) and the setting based on the number of SNPs within the genomic region ((p1=p2=1/(number of neighbor SNPs within the region), p12=p1/10). Traits are considered colocalized if either parameter configuration produces a posterior probability for shared causality (PP4) exceeding 0.7.

## Code availability

All code and results will be made available at https://github.com/jke20/BRONTE-BrainTissueGNN.git

## Result

### Overview of Brain Ontology Expression (BRONTE) model

Our model BRONTE expands the GTEx transcriptomic data by integrating it with the Allen Brain Atlas of gene expression, a resource which comprehensively assays gene expression across human brain tissues.The model is trained on 10 out of 13 GTEx brain tissues, excluding two duplicate tissues, and one tissue absent in the Allen dataset. From the Allen Brain Atlas, we focus on 103 tissues within the left hemisphere shared across 6 subjects. We model expression of 15,044 genes present in both datasets.

BRONTE’s design is based on the concept that expression of a particular gene, in a particular brain tissue, for a particular subject can be predicted based on integrating representations of gene expression variation for each of the tissue, the gene, and the subject. To this end, BRONTE’s input uses the normalized expression values for a subject for that gene, across the 10 GTEx tissues, to represent the subject, along with learned tissue and gene embeddings, outlined below. These representations comprise input to a neural network trained to predict expression of that gene, for that subject, for all 103 Allen brain tissues, including the 10 GTEx tissues.

BRONTE’s design leverages graph neural networks (GNNs), which learn similar embeddings for neighboring nodes. Here, each node is one brain tissue, and the graph structure is derived from the Allen brain atlas ontology, which represents the hierarchical organization of brain tissues. The Allen brain project demonstrated that functionally related tissues exhibit similar gene expression patterns, and we expect our GNN to capture these relationships among brain tissues. In addition to the tissue representation, the gene embeddings are trained to represent variation in expression of that gene across tissues, based on the finding from the Allen brain project that genes functioning in brain tissue display replicable tissue-specific expression patterns. Our model is trained on both Allen and GTEx subjects, as outlined in the Methods.

### Imputed gene expression recovers variation among subjects, genes, and tissues

To evaluate the performance of BRONTE, we first assess its ability to predict gene expression for GTEx subjects in uncollected tissues, both within the original 10 GTEx brain tissues, and expanding beyond those 10 tissues to 103 Allen brain tissues.

Initially, we focus on the imputation of gene expression within the 10 GTEx brain tissues, by assessing performance in held-out expression samples for a subject. Only 30 of 300 GTEx subjects have gene expression for all 10 tissues; BRONTE allows us to effectively expand the GTEx dataset to include the uncollected tissues. Because the primary use of GTEx is to identify genetics influencing expression of a gene within a tissue, we focus on predicting relative expression of a gene across individuals, within a brain tissue. We compare held out gene expression to predicted values. We evaluate BRONTE’s predictive performance against four alternative models—linear regression model (LM), MICE[15], and CP method[16], all of which use the gene expression data to predict, as well as PrediXcan[17], which predicts expression based on genotype (see Methods). For each brain tissue, we calculate a Spearman correlation coefficient for each gene by comparing the predicted and held-out gene expression across GTEx subjects, and we compare the distribution of Spearman correlation coefficients within a tissue, between different models. We also demonstrate the robustness of our model and its ability to expand the GTEx dataset by imputing data for GTEx subjects up to five missing tissues (Supplementary Figure 1). BRONTE performs at least as well as all other models in nine out of ten tissues, with LM outperforming BRONTE in one tissue (Hippocampus), and CP method, MICE, and PrediXcan showing consistently inferior performance.

**Figure 1.**
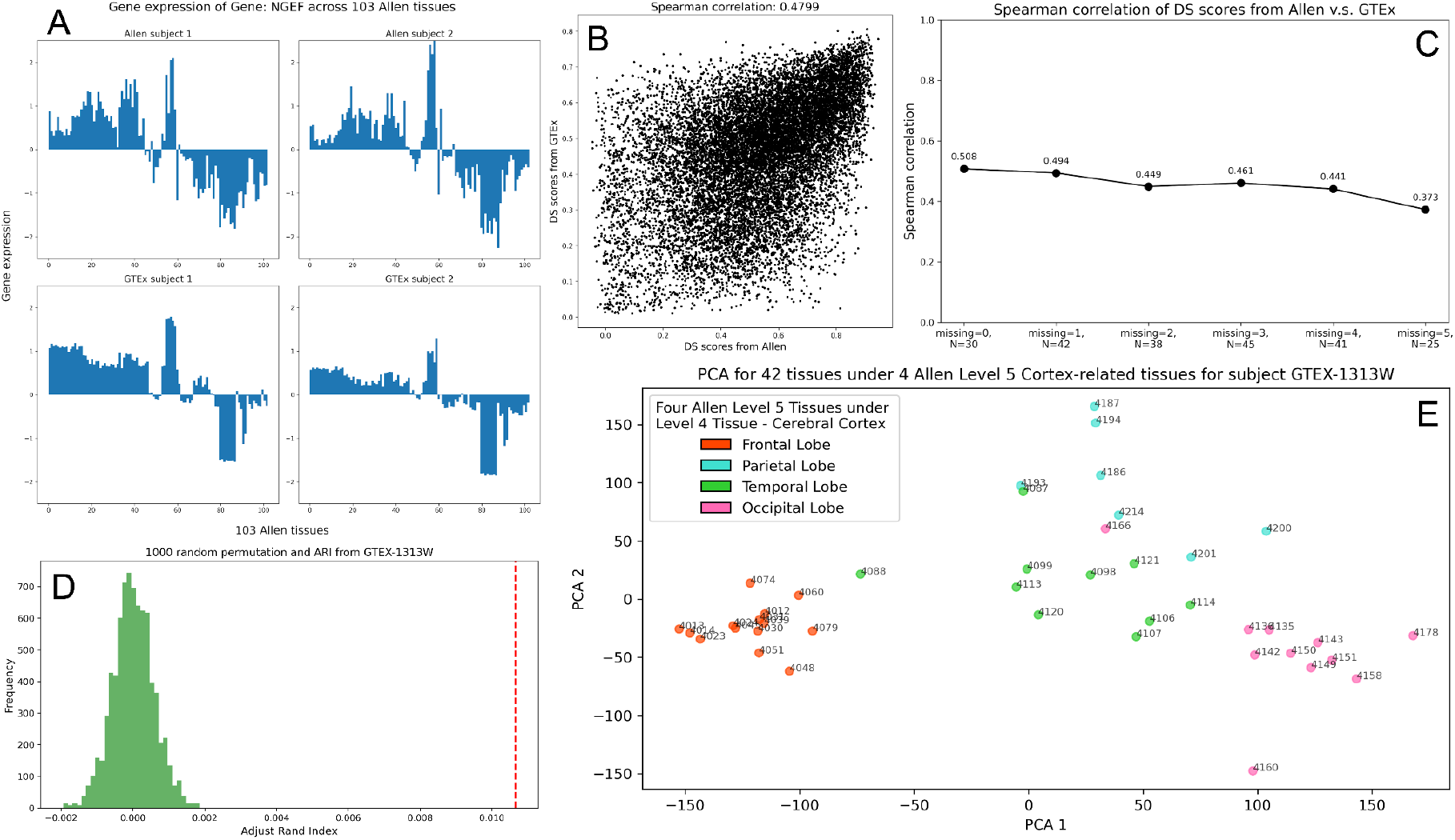
Evaluation of BRONTE’s prediction. **A**, Bar ptots showing the gene expression of a gene with high DS score, NGEF, across 103 Allen Issues for two Allen subjects (top) and two GTEx subjects (bottom). B. Comparison between the DS scores from Allen dataset and those from our results on the overtapped genes. Each dot represents one gene, comparing DS score In Allen brain (x-axis) against DS score in GTEx imputed data (y-axis). Genes with higher DS scores In Allen also have higher DS scores In the imputed data for all 221 GTEx subjects (Spearman correlation = 0 46) **C**, Spearman correlation between DS scores from the Allen dataset and our results derived from GTEx subjects with varying numbers of missing tissues Here, ‘missing’ refers to the number of missing tissues, and **‘N’** indicates the number of GTEx subjects with that exact number of missing tissues. **D**, Comparing the ARI from our WGCNA-denved modules (red line on the right) with 1000 random ARIs from permutations (distribution in green) using the gene expression of subject GTEx-1313W. **E**, Pnncipal Component Analysis on 42 Allen tissues under 4 Allen Level 5 Cortex-related tissues for subject GTEx-1313W.

Next, we evaluate BRONTE’s ability to expand beyond the GTEx tissues, predicting gene expression across 103 Allen brain tissues for GTEx subjects. The competing predictive models we used to evaluate GTEx predictions are not able to expand beyond the GTEx tissues, a key distinction of BRONTE. Instead, we evaluate whether our prediction accurately reflects biologically meaningful expression patterns previously reported in Allen tissues. Previous work[18] used the Differential Stability (DS) scores to quantify how stable a gene’s brain tissue expression patterns are between subjects: for example, Figure 1A shows a gene with high DS score across 103 Allen tissues for two Allen subjects and two GTEx subjects. For each gene, we calculate DS scores using the imputed Allen tissues for GTEx subjects with at least five GTEx brain tissues and we compare these scores to those from the Allen dataset using Spearman correlation (Figure 1B). The results show that genes with high DS scores in the Allen dataset exhibit high DS scores in the GTEx predictions (Spearman correlation=0.48), indicating strong alignment between our predicted and observed expression patterns across different Allen brain tissues.

Since BRONTE predicts gene expression for GTEx subjects with varying levels of data completeness, we also evaluate its robustness with missing data by repeating the DS score analysis for GTEx subjects with up to five missing tissues (Figure 1C). We observe stable model performance, even when subjects have up to five missing tissues.

Previous analysis of the Allen Brain Expression Atlas identified clusters of genes associated with distinct biological functions using Weighted Gene Co-expression Network Analysis (WGCNA)[19]. To assess whether our predictions recapitulate these gene expression patterns, we apply WGCNA to the gene expression data from 103 predicted Allen tissues across GTEx subjects, and we compare our WGCNA-derived modules with the original Allen modules, calculating the Adjusted Rand Index (ARI, see Methods). By benchmarking the score against ARI values from 1,000 random permutations, we show that the ARI from our results is significantly elevated for subject GTEX-1313W (p-value < 1e-3, see Methods, Fig 1D). Similar trends are observed across other GTEx subjects with at least five tissues. These findings indicate that BRONTE’s imputed expression for GTEx subjects in Allen tissues preserves the clustering patterns of the original Allen data, demonstrating that biologically meaningful signal is retained.

Finally, we assess whether our predictions capture the spatial organization of gene expression patterns observed in the original Allen study. In that study, Principal Component Analysis (PCA) of the expression data demonstrated that sub-tissues within the same major cortical tissue have similar gene expression profiles. We conduct a PCA analysis on the imputed gene expression of subject GTEX-1313W across 42 Allen tissues within the four major cortical subdivisions of the Cortex: Frontal Lobe, Parietal Lobe, Temporal Lobe, and Occipital Lobe. Our PCA results show that genome-wide gene expression profiles are most similar for tissues within the same lobes, indicating that BRONTE accurately reflects both inter- and intra-tissue expression patterns (Figure 1E).

Taken together, these analyses show that our predictions successfully recover the biological variation among subjects, genes, and tissues.

### Expanded GTEx data improves discovery of eQTLs

The GTEx project was originally designed to enable the identification of expression quantitative trait loci (eQTLs), providing a public dataset of genetic variants associated with gene expression. We perform eQTL analysis using the GTEx genotype data and matched generated expression data on 221 people with at least five measured brain expression profiles. To verify that our findings on the expanded GTEx dataset align with the original GTEx findings, we compare the p-values and effect sizes of overlapping eQTLs from the original GTEx data and our expanded dataset. The results for one of the 10 GTEx tissues, the Amygdala, are shown in Figure 2A and 2B. Other tissues have similar results, shown in Supplementary Figure 2, 3. Our analysis with the expanded data generally yielded smaller p-values, indicating increased significance for many previously identified eQTLs. The strong correlation between both p-values (Spearman correlation = 0.84) and effect sizes (Spearman correlation = 0.99) suggests that our expanded dataset closely replicates the original GTEx results while enhancing the statistical power of eQTL associations.

**Figure 2.**
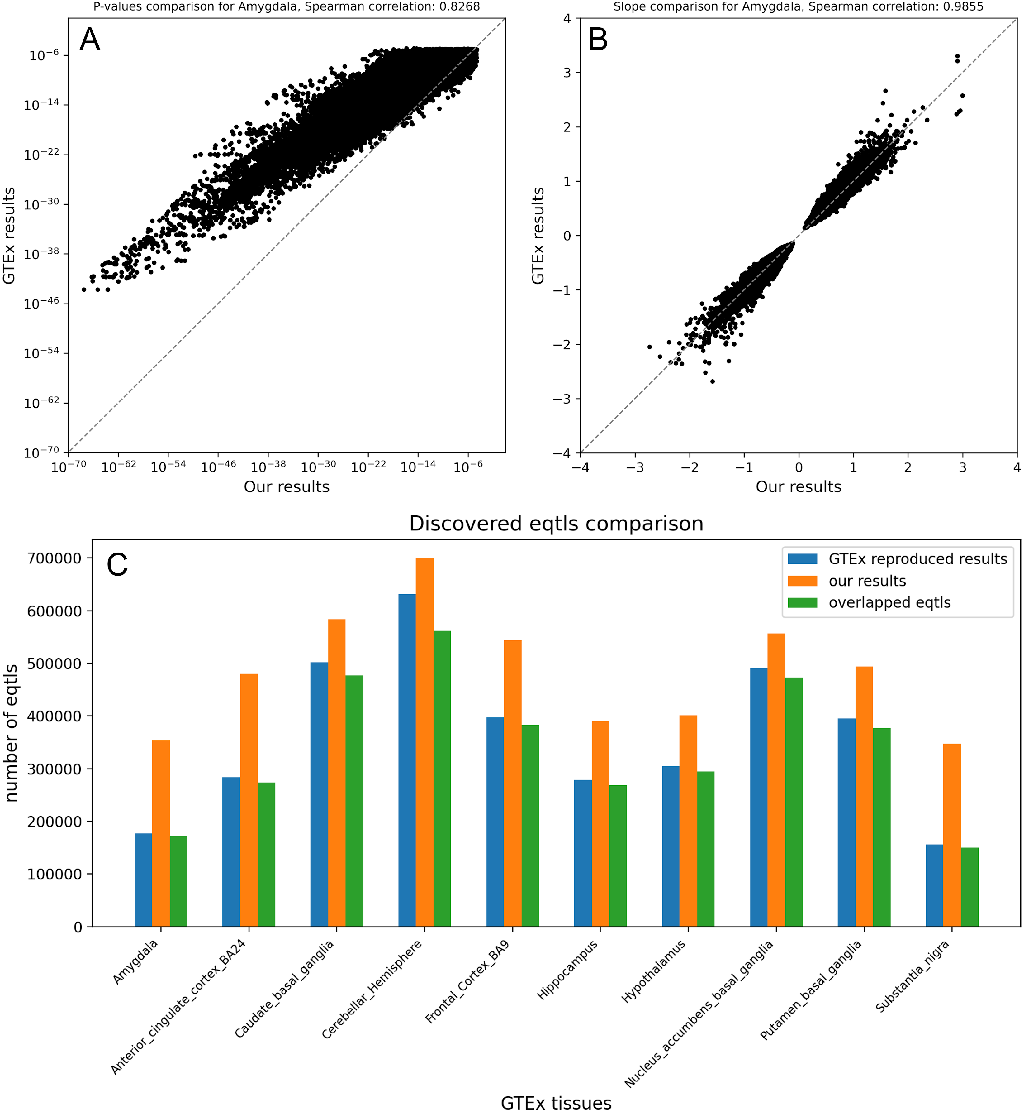
eQTL analysis on 10 GTEx tissues. **A.B**, P-value **(A)** and Slope **(B)** comparison between our results and GTEx results on the overlapped eQTLs for GTEx tissue Amygdala-Each point is one eQTL. **C**, Bar plots showing the number of eQTLs identified in three categories: GTEx expression data eQTLs. BRONTE imputed data eQTLs. and the overlapping eQTLs across 10 GTEx tissues. “GTEx reproduced results” refers to eQTLs obtained by running our eQTL analysis on the original GTEx data, rather than directly using results from the GTEx portal, to ensure consistency.

**Figure 3.**
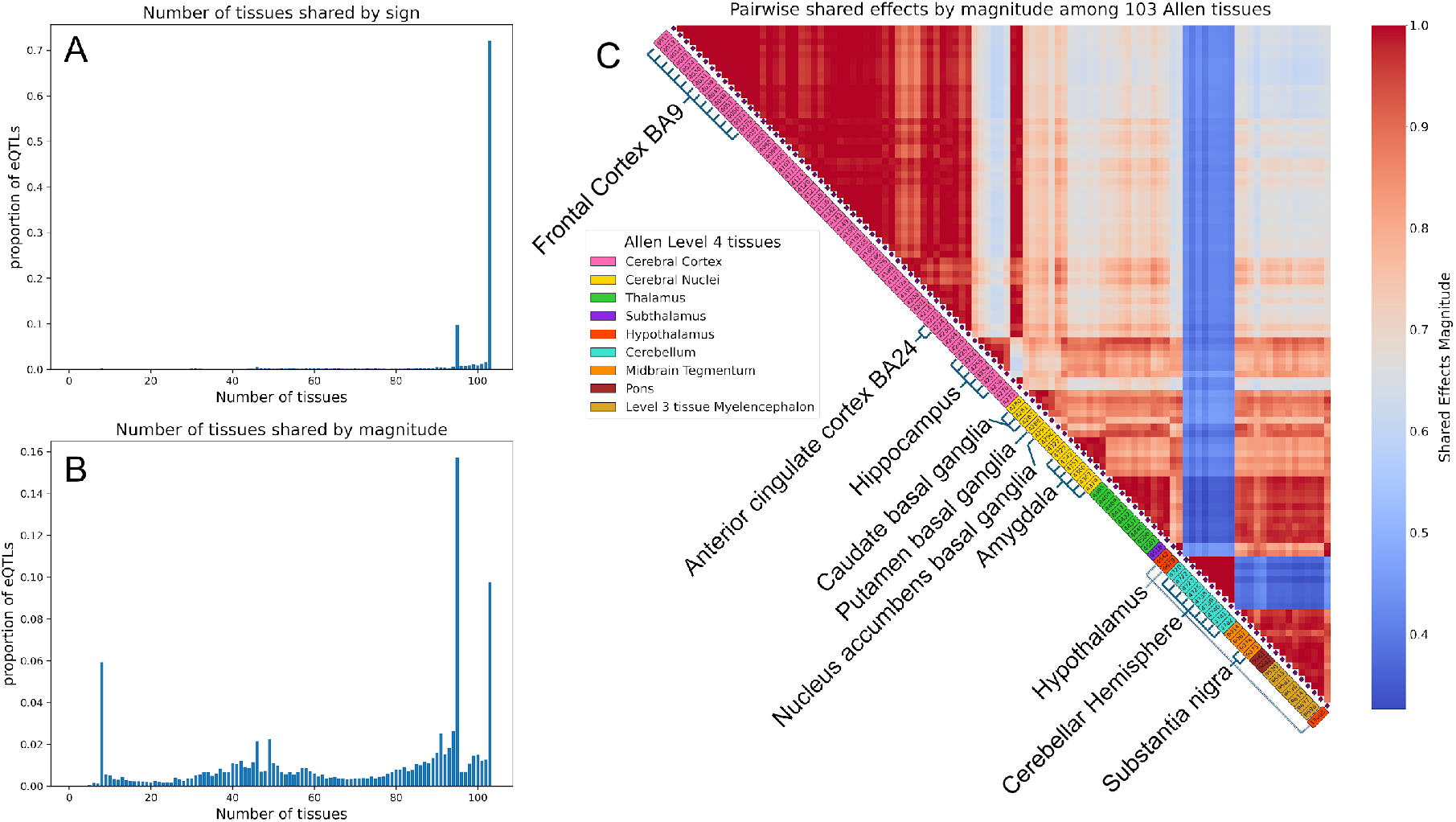
eQTL analysis on 103 Allen tissues. **A**,**B**, Histograms showing estimated number of tissues in which lead eQTLs are “shared”, using two different sharing definitions, by sign **(A)** and by magnitude **(B). C**, Pairwise sharing of eQTL effects by magnitude across 103 Allen tissues, with tissues grouped by Allen Level 4 categories and distinguished by color. The mapping between the 10 GTEx tissues and the Allen tissues is illustrated in the accompanying diagram.

We formally assess the overlap in results from the original and imputed data in Figure 2C. This result shows that our expanded dataset enables the discovery of new eQTLs while successfully reproducing most previously identified ones.

### eQTL analysis on 103 Allen brain tissues for GTEx subjects

We expand the GTEx data to 103 tissues and perform eQTL discovery, and then we seek to assess whether these effect estimates are plausible by comparing them to previous brain eQTL analyses. In particular, Urbut et al. (2019) investigated sharing of eQTL effect estimates among pairs of brain tissues, with two metrics: sharing of effects by sign (direction of effect) and sharing by magnitude. Sharing by magnitude refers to effects that share signs and differ by no more than a factor of two in size. Their findings confirmed extensive eQTL sharing among brain tissues, with up to 96% shared by sign and 76% shared by magnitude[26].

We use this result to evaluate the eQTL effects discovered in our imputed data. That is, we expect that a high level of shared effects will indicate that our predictions successfully capture the extensive sharing of eQTL effects among brain tissues observed by Urbut et al. For each lead SNP with strongest eQTL signal in a region of linkage disequilibrium, we examine sharing by both sign and magnitude, across 103 Allen tissues. Our results show that 72.0% of lead eQTLs have shared effects by sign across 103 Allen tissues (Figure 3A), consistent with Urbut’s work. In Figure 3B, most lead eQTLs also show shared effects by magnitude (peaks at the right side of the histogram). In both types of sharing, the Cerebellar Hemisphere (represented by eight more refined Allen brain tissues) is the brain tissue largely inconsistent with the other tissues, a signal also found in the work by Urbut.

Intuitively, Allen tissues within the same major anatomical area should show similar eQTL sharing effects, reflecting anatomical and functional coherence. To test this, we examined pairwise sharing of eQTL effects by magnitude across the 103 Allen tissues (Figure 3C). Our results confirm that Allen tissues within the same GTEx group and Allen major anatomical areas show higher pairwise sharing, with Cerebellar Hemisphere-related tissues displaying strong within-group sharing but less sharing with other brain tissues. This overall pattern highlights anatomical and functional specificity in eQTL sharing, supporting the accuracy of BRONTE in capturing biologically meaningful relationships.

### Discovery of genes and tissues involved in brain-related traits

As a major goal of eQTL analysis is discovery of the functional effects of genetic variation, we apply Mendelian Randomization (MR) analysis to assess causal relationships between eQTL-mediated gene regulation and brain-related traits. We test 11,841 significant eQTLs across 103 Allen brain tissues, and 30 brain-related traits (Supplementary Table 2), finding that 4,734 MR results remain significant after false discovery rate correction (see Methods). Then, we perform colocalization analysis to determine whether the eQTLs and brain-related traits share a common genetic signal. Of the MR-significant eQTLs, 2,955 also show evidence of co-localization, including 150 genes (Figure 4A, B). Summarized results for these 2,955 eQTLs are provided in Supplementary Table 3.

**Table 2.** Testing trait association with brain regions.

| Trait | Allen Level 4 Tissue | P-value |
| --- | --- | --- |
| MDD | Cerebral Cortex | 1.10e-10 |
| BD | Cerebellum | 5.38e-07 |
| Generalized EPI 2 | Cerebral Nuclei | 1.27e-05 |
| ADHD | Cerebral Nuclei | 1.22e-04 |
| ALS | Cerebral Cortex | 2.52e-03 |
| Schizophrenia | Cerebellum | 6.55e-03 |
| Cognitive function | Hypothalamus | 7.34e-03 |
| Putamen volume | Midbrain Tegmentum | 3.32e-02 |
| Years of schooling | Thalamus | 3.83e-02 |

**Table 3.**
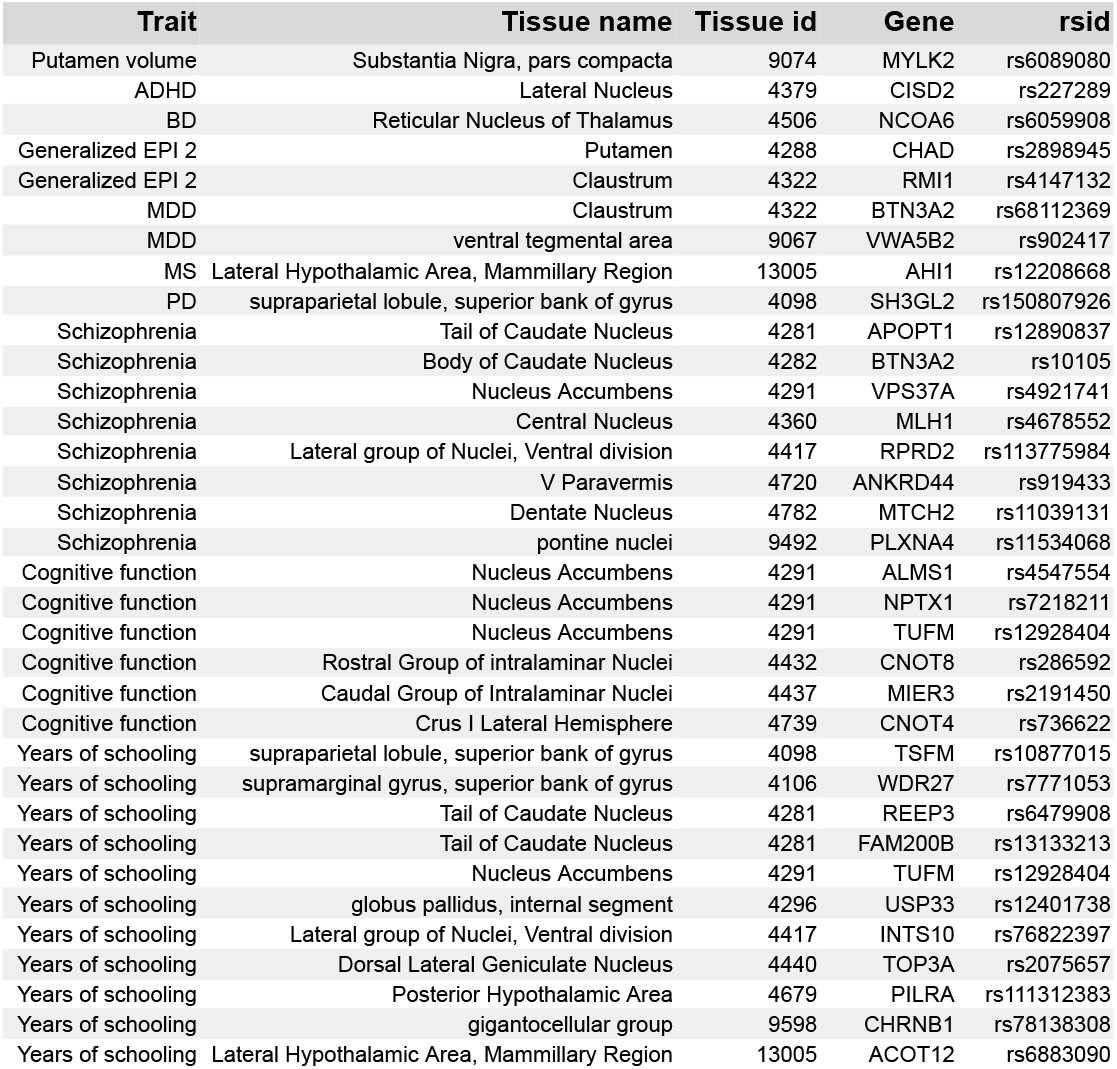
Significant tissue-specific SNPs in Traits.

**Figure 4.**
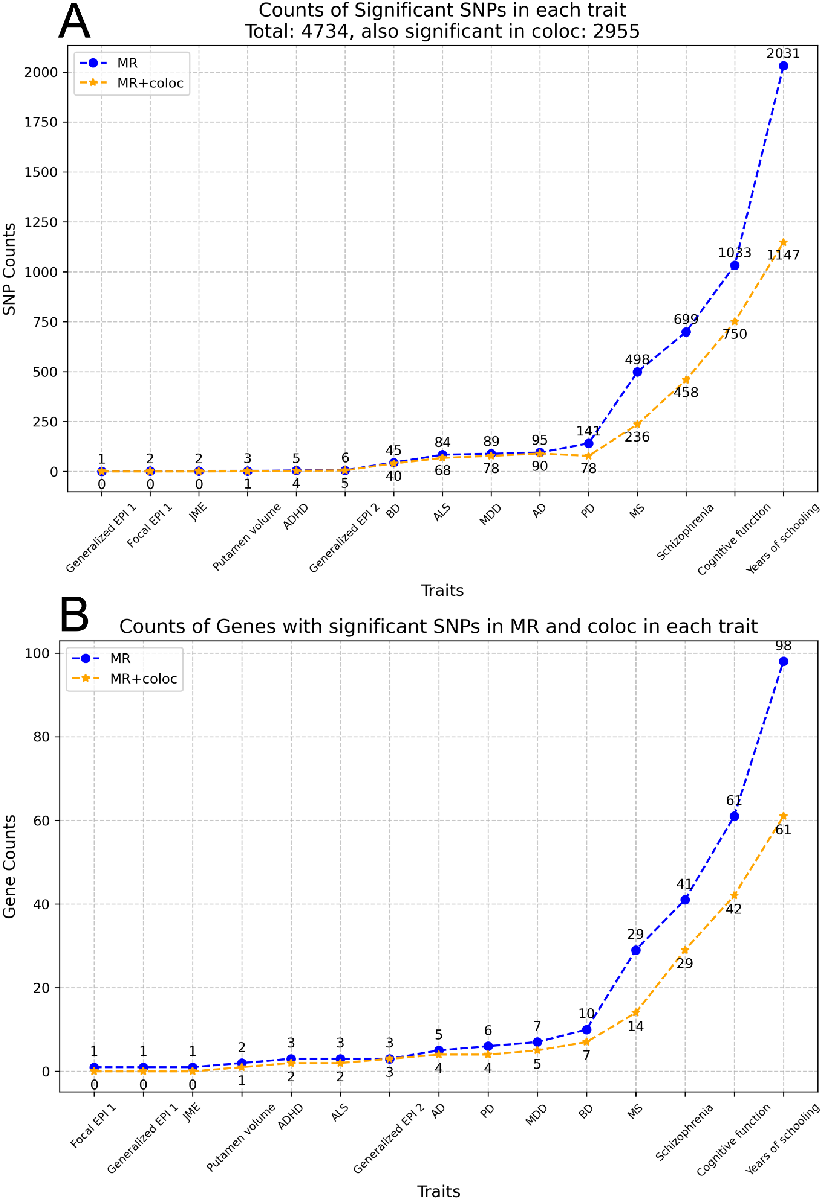
MR analysis and Colocalization for brain-related traits. A.B. Counts of SNPs **(A)** and Counts of Genes with SNPs **(B)** that are significant in the MR analysis (blue dots) and in both the MR analysis and colocalizalion (yellow dots) for brain-related traits, sorted in the ascending order. Full names of traits are provided in Supplementary Table 2.

Next, we assess whether our gene-level discoveries replicate those reported in the Metabrain study[27], as shown in Table 1, focusing on 11,561 genes shared between the two datasets. Table 1 summarizes the overlap of BRONTE’s results with these findings, demonstrating significant overlap with those reported in the Metabrain study across eight out of thirteen traits.

One of BRONTE’s goals is to enable discovery of the tissue where changes in regulation of a gene can impact brain-related traits. In order to identify the important tissues for each brain-related trait, we first visualize the number of eQTLs that are significant in both MR analysis and colocalization for each trait (Supplementary Figure 4). To quantitatively assess whether specific traits are statistically associated with particular broad brain regions, we test enrichment of causal eQTLs for each trait within each broad (Allen Level 4) region, as compared to the other brain related traits, and compared to the other tissues. The significantly enriched broad regions for each brain-related trait is summarized in Table 2. These results pinpoint potential causal brain areas that may drive brain-related traits. To identify genes with tissue-specific effects, we examine 2,955 SNPs that are significant in both MR analysis and colocalization. Among these, we identify 34 SNPs associated with genes that exhibit significance exclusively in a single Allen brain tissue, without significant associations in any other tissue (Table 3). For instance, we recover a novel association of ADHD with expression of *CISD2*, in the lateral nucleus, a structure within the amygdala. Reduced amygdala volumes have been observed in patients with ADHD[28], and recent proteome-wide association studies have identified *CISD2* as one of several proteins significantly linked to ADHD[29]. However, the specific pathogenic tissue within the Amygdala and the pathways underlying ADHD development remain poorly understood. This is just one example of how BRONTE is able to reveal a unique subset of genes that may drive brain tissue-specific processes, providing critical targets for advancing our understanding of brain-related traits.

## Discussion

We put forward a novel model BRONTE that imputes the relative expression across subjects in a given gene and tissue, in order to recover the association of genetics with tissue-specific gene regulation. Our model is based on data showing that gene expression follows characteristic patterns across brain tissues, making it the first model to employ the hierarchical relationship of brain tissues to impute brain expression. We demonstrate that BRONTE’s gene expression predictions recover known variation across genes and brain tissues. Most importantly, our rigorous analysis demonstrates that the imputations also recover variation across subjects, within a single gene and tissue. BRONTE’s imputations allow recovery of brain eQTL effects that validate against published eQTLs. As well, we discover causal effects of gene regulation on disease that significantly reproduce those previously reported.

Both the increased sample size, and the increased tissue diversity imputed by BRONTE improve discovery of genes that may impact brain traits. Higher sample size enables discovery of eQTL effects; by increasing the number of samples from 165 to 210 on average per tissue, our imputed data set is able to discover from 11.0 to 122.9% more eQTLs per brain tissue. But our particular focus is on improving the diversity of tissues available for each brain sample, enabling discovery of novel disease driver genes that may exert their effect in overlooked tissues. For example, the Metabrain analysis integrating eQTLs and GWAS variants did not discover any genes for ADHD or generalized epilepsy. In contrast, our ADHD finding is described above, while for epilepsy we recovered gene *RMI1*, recently also highlighted by a larger GWAS meta-analysis of epilepsy[30].

The diversity of brain tissues also allows us to put forward intriguing new hypotheses about the brain regions involved in development of brain-related traits. Some findings are supported by recent literature: the cerebral cortex and cerebellum have been implicated in the pathophysiology of major depressive disorder (MDD) and bipolar disorder (BD), respectively. In MDD, structural MRI studies reveal cortical thinning in the prefrontal cortex, a tissue within the cerebral cortex critical for emotional regulation and cognitive processing[31]. Functional and structural abnormalities in the prefrontal cortex are further linked to core depressive phenotypes, such as negative processing biases and anhedonia, underscoring its pivotal role in MDD[32]. In BD, elevated T1ρ values in the cerebellum indicate potential metabolic or microstructural disruptions, emphasizing the cerebellum’s contribution to the disorder[33]. Additionally, altered cerebellar tract integrity has been associated with different stages of BD, highlighting its relevance to the progression of the disease[34].

BRONTE does have a number of limitations: we are only able to impute expression for genes shared between Allen brain and GTEx. As well, there is no easy way to validate the expression for uncollected GTEx tissues, meaning we must rely on indirect means of evaluating the model’s performance. Our MR and colocalization results do not recover all Metabrain causal genes, likely because their focus was on improving the sample size in the cortex.

Because direct assays of human brain tissue are limited and expensive, BRONTE promises to amplify the discoveries possible from existing data sets. Our model, the imputed gene expression data across 103 tissues, and our discoveries for therapeutic targets, are openly available resources to improve understanding of mechanisms of effect for variants linked to brain related traits.

## Funding sources and Grant numbers

This work was supported by the National Institute of General Medicine Sciences (NIGMS R35 GM151001-01) to RDM.

