## Supplementary figures and images for "Discovering disease genetic variation impacting gene expression in 103 brain tissues with the Brain Ontology Expression (BRONTE) graph neural network model"

### Supplementary Figure 1.docx

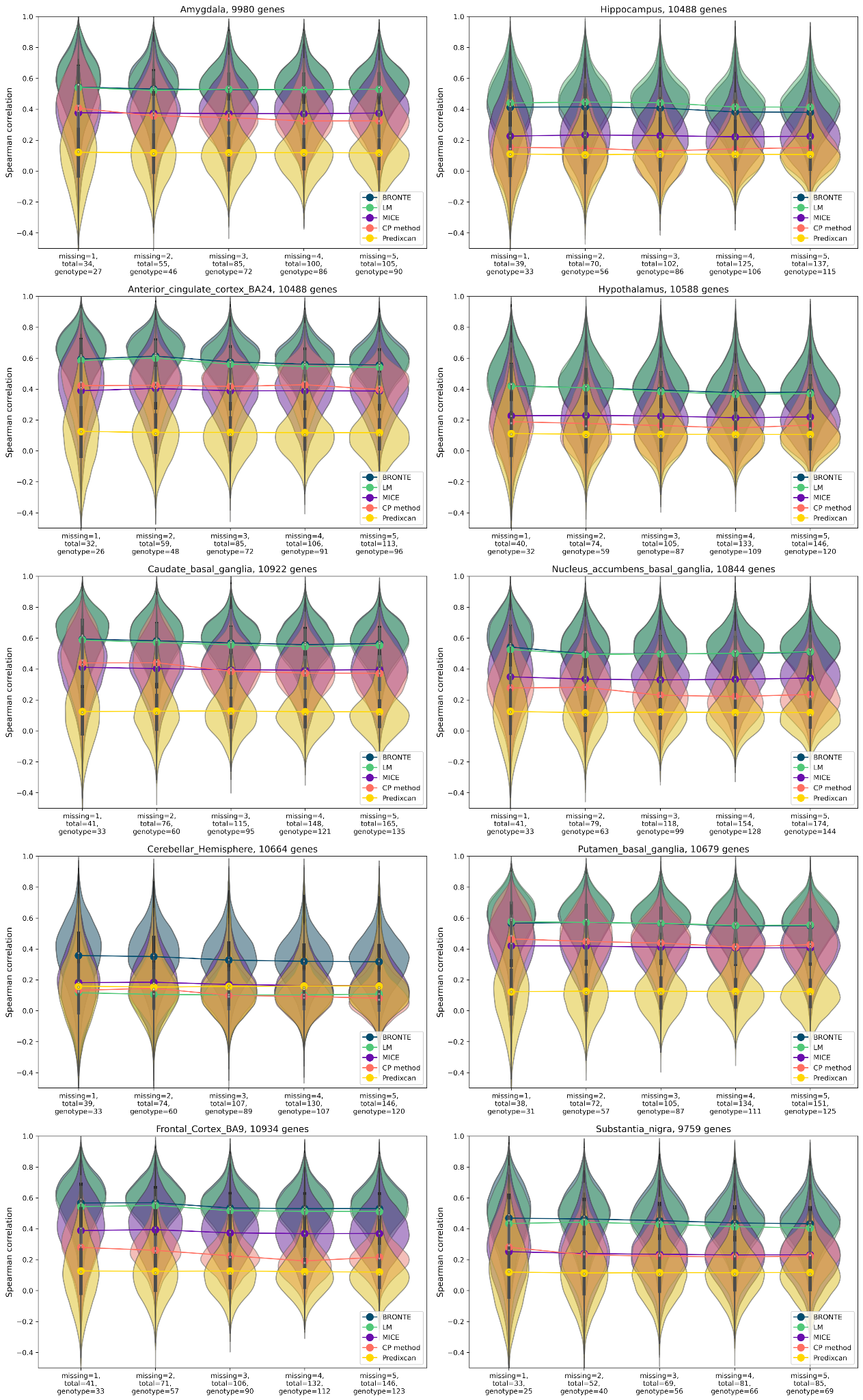

### Supplementary Figure 2&3.docx

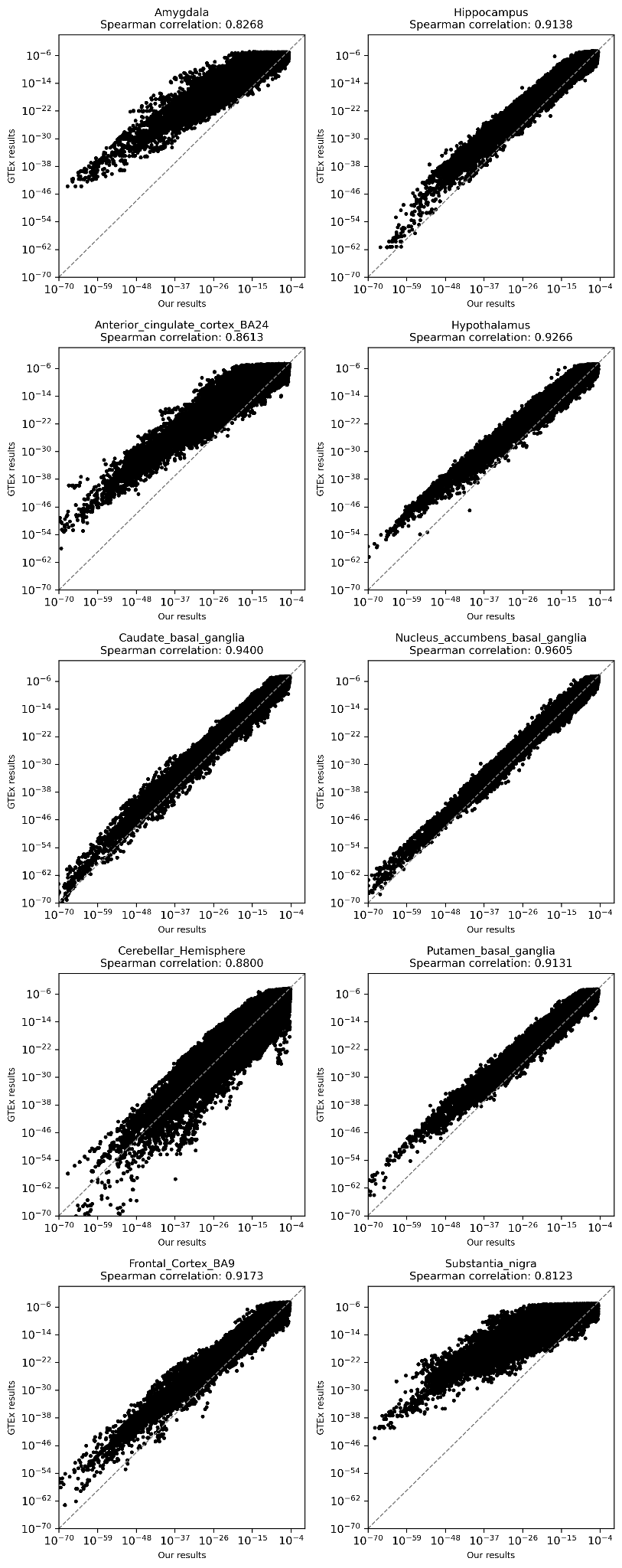

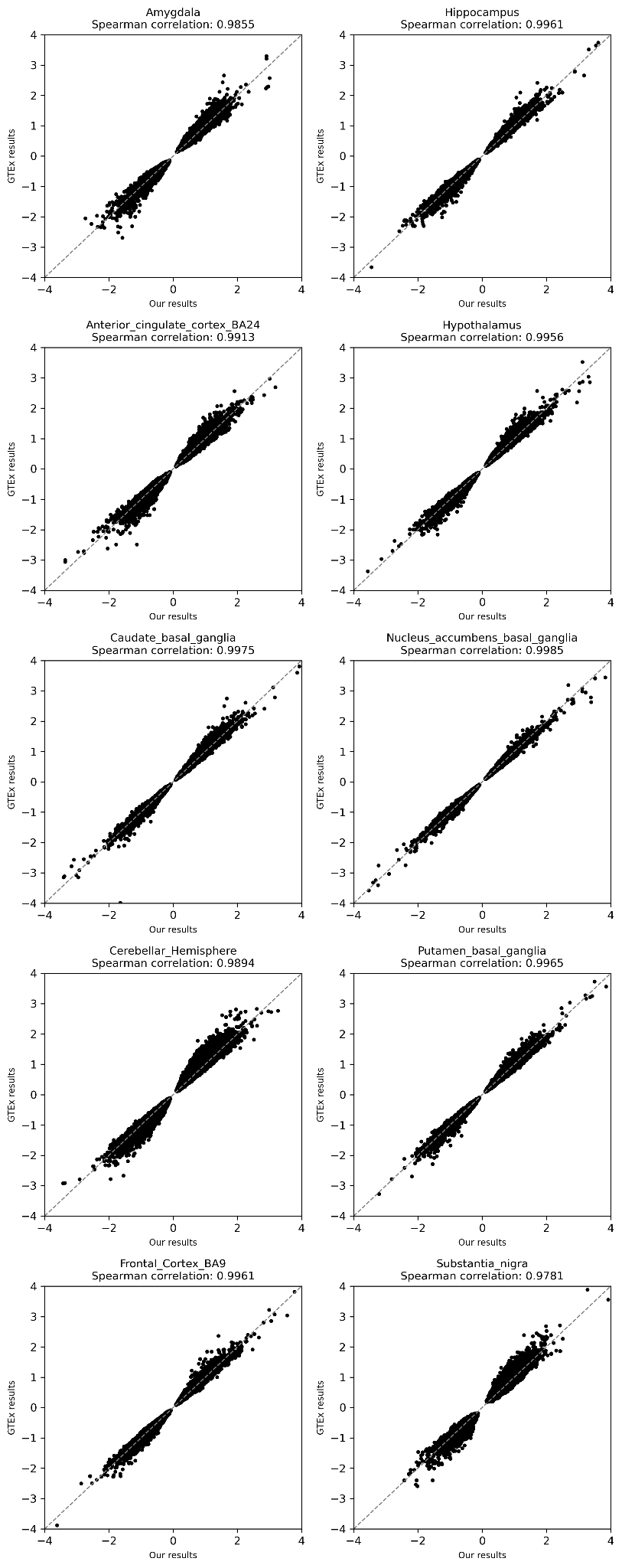

### Supplementary Figure 4.docx

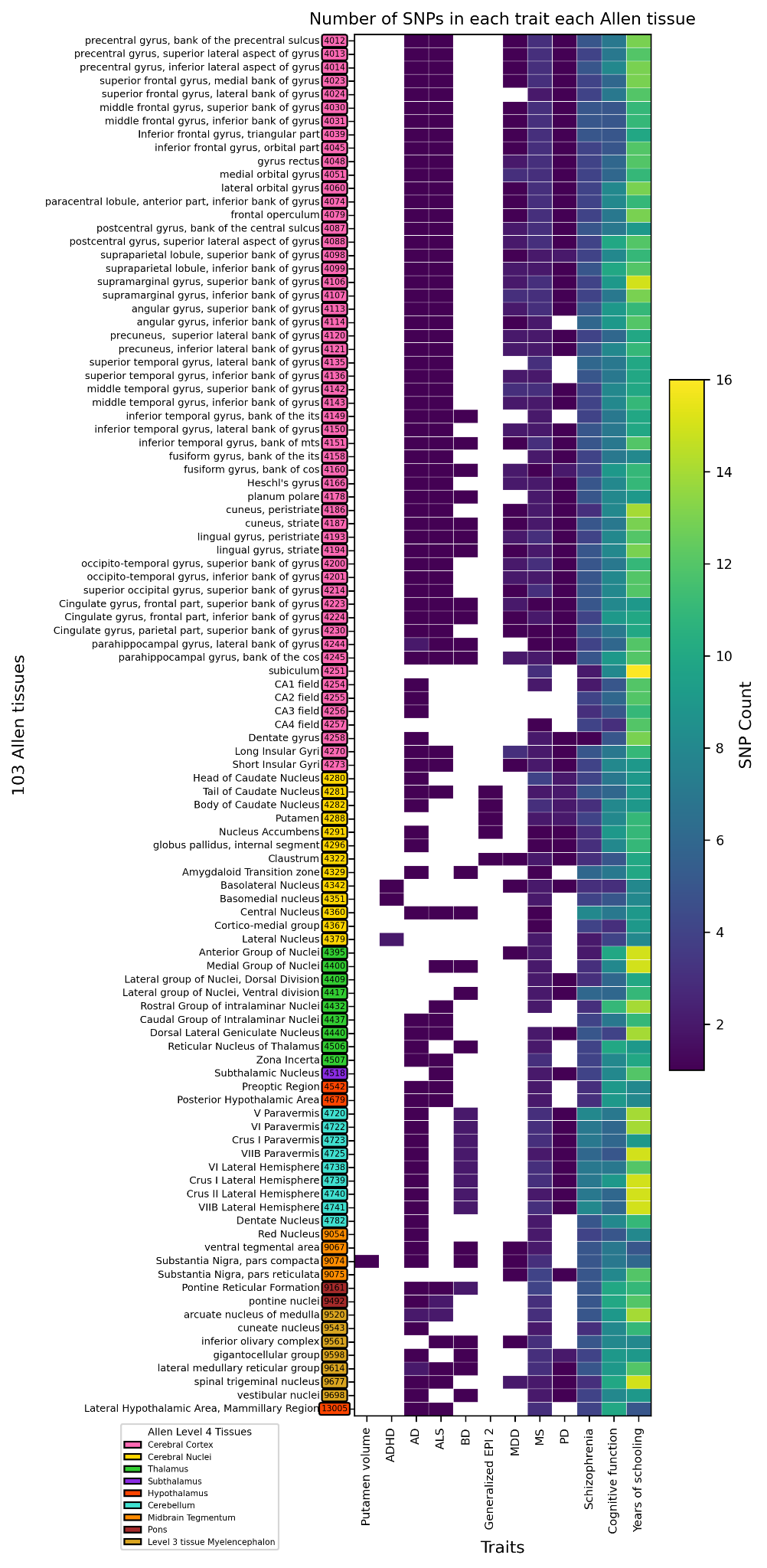
